# REL2 overexpression in the *Anopheles gambiae* midgut causes major transcriptional changes but fails to induce an immune response

**DOI:** 10.1101/2024.02.05.578852

**Authors:** Astrid Hoermann, Paolo Capriotti, Giuseppe Del Corsano, Maria Grazia Inghilterra, Tibebu Habtewold, Julia A. Cai, Gauri Sachiko Saini, Huong Nguyen, Nikolai Windbichler, George K. Christophides

## Abstract

The NF-κB-like transcription factor, REL2, is a key player in the mosquito Immunodeficiency (Imd) pathway and holds promise for controlling malaria parasite infections in genetically modified *Anopheles gambiae* mosquitoes. We engineered transgenic mosquitoes overexpressing REL2 from within the bloodmeal-inducible zinc carboxypeptidase A1 (CP) host gene in the adult posterior midgut. Our results confirmed elevated REL2 expression in the posterior midgut following a bloodmeal, with the corresponding protein localized within epithelial cell nuclei. While this induced overexpression triggered substantial transcriptional changes, accompanied by notable fitness costs, the resultant reduction in *Plasmodium falciparum* infection was modest. An in-depth analysis of regulatory regions of differentially regulated genes allowed us to identify direct REL2 target genes and revealed signatures indicative of potential transcriptional repressors. To account for potential impacts of host gene modification, we also created a CP knockout line that caused marginal effects on mosquito fitness. These findings shed light on the observed absence of transcriptional activation and, in some cases, induced repression of antimicrobial peptides (AMPs) presumed to be under Imd pathway control. In conclusion, our study suggests that elevated REL2 expression in the posterior midgut may induce the upregulation of negative immune regulators, facilitating control over an otherwise unrestrained immune response, and that concurrent transcriptional derepression may be needed to effectively induce the mosquito immune response. This work contributes valuable insights into the intricate regulation of midgut immunity in malaria vector mosquitoes.

## Introduction

REL2 is a key regulator in the mosquito immune response against bacteria and malaria parasites, orthologous to the *Drosophila* NF-κB-like transcription factor, Relish. Receptor-mediated recognition of bacteria and other pathogens activates the Imd pathway that through REL2 causes transcriptional induction of AMPs and other immune effectors [1]. The full-length isoform of REL2, REL2-F, features carboxyl-terminal ankyrin (ANK) repeats and a death domain (DD), together concealing the nuclear localization signal (NLS) [2]. Proteolytic cleavage of these inhibitory regions frees the amino-terminal Rel-homology domain (RHD), prompting REL2 translocation to the nucleus. A shorter isoform, REL2-S, lacking the ANK repeats and DD, has also been identified [2], but its physiological role remains unclear.

Silencing Caspar, a negative regulator of the Imd pathway, is shown to significantly impede mosquito infection with the human malaria parasite *P. falciparum* across various *Anopheles* species [3]. Conversely, ectopic REL2 expression in the fat body of *Aedes aegypti*, controlled by the bloodmeal-inducible vitellogenin promoter, activates the expression of defensins and cecropins, leading to reduction in susceptibility to the avian malaria parasite *P. gallinaceum* [4]. Transcriptomic analysis 24 hours (h) post bloodmeal in this transgenic *Aedes* strain revealed that over half of the induced genes function in immunity. A hybrid strain, combining overexpression of REL2 with that of REL1, the Toll pathway transcription factor, co-orthologous to *Drosophila* Dorsal and Dif, demonstrated a synergistic action between these two transcription factors [5].

*An. gambiae* REL2-S overexpression in transgenic *An. stephensi*, driven by the *An. gambiae* CP promoter, is also shown to significantly reduce *P. falciparum* infection [6]. Transcriptome and proteome analysis of this CpREL2_15_ transgenic line revealed that only a small number of differentially expressed genes (DEGs) belonged to the immunity functional category [7]. The authors suggested that potential targets of the REL2-mediated antimalarial effect in this *An. stephensi* line included alpha-2-macroglobulin (A2MRAP), leucine-rich transmembrane protein (LRTP), REL2-responsive serine protease 2 (R2RSP2), angiotensin converting enzyme precursor (ACEP), and serine protease precursor 2 (SRPN2). Furthermore, the line exhibited mating preferences, with transgenic males favoring wild-type (WT) females, and *vice versa*, an effect attributed to their reduced microbial load [8].

Building upon these prior studies, we sought to explore whether inducing endogenous REL2-S overexpression in the *An. gambiae* midgut post-bloodmeal could serve as a viable strategy for modifying *An. gambiae* populations toward malaria elimination, when combined with gene drive. To achieve this, we employed the Integral Gene Drive (IGD) conceptual framework [9], previously proposed for *An. gambiae* population replacement, by integrating REL2-S and an intronic guide RNA (gRNA) module within the somatic CP locus. This modification could be driven through populations via a Cas9 endonuclease expressed in the germline and integrated into a distinct genomic locus. Our earlier work has already demonstrated the successful modification of the *An. gambiae* CP locus (along with other midgut loci) to accommodate such antimalarial effectors and their corresponding gRNA [10, 11]. These modifications demonstrated high non-autonomous homing rates and minimal fitness costs, highlighting the potential of IGD as a powerful tool for driving the expression of antimalarial effectors in mosquito populations. Contrary to our expectation, the results suggest that the sole overexpression of REL2 in the midgut is not an effective strategy for *An. gambiae* population modification, as it fails to induce a strong immune response and has modest impact on malaria infection. Instead, REL2 triggers the expression of several transcriptional repressors among many other transcriptional changes, which may contribute to an observed immune paucity and notable fitness costs.

## Results

### Generation of transgenic *An. gambiae* overexpressing REL2-S from within the CP locus

We previously evaluated CP (AGAP009593) for its suitability as a host locus for effector integration, including its mRNA expression with and without a fluorescent marker module, as well as its homing potential through an intronic CP gRNA [11]. The same insertion site was used here for a 5’ in-frame insertion of REL2-S after the CP start codon (**Figure 1A**). The P2A cleavage peptide was used to fuse the effector cassette to the CP host gene, which contains a secretion signal. An intron was inserted within the REL2-S coding sequence (CDS), harboring the U6-gRNA module required for homing and subsequent gene drive as well as the green fluorescent protein (GFP) expression under the control of the synthetic 3xP3 promoter required for transgenesis.

**Figure 1.**
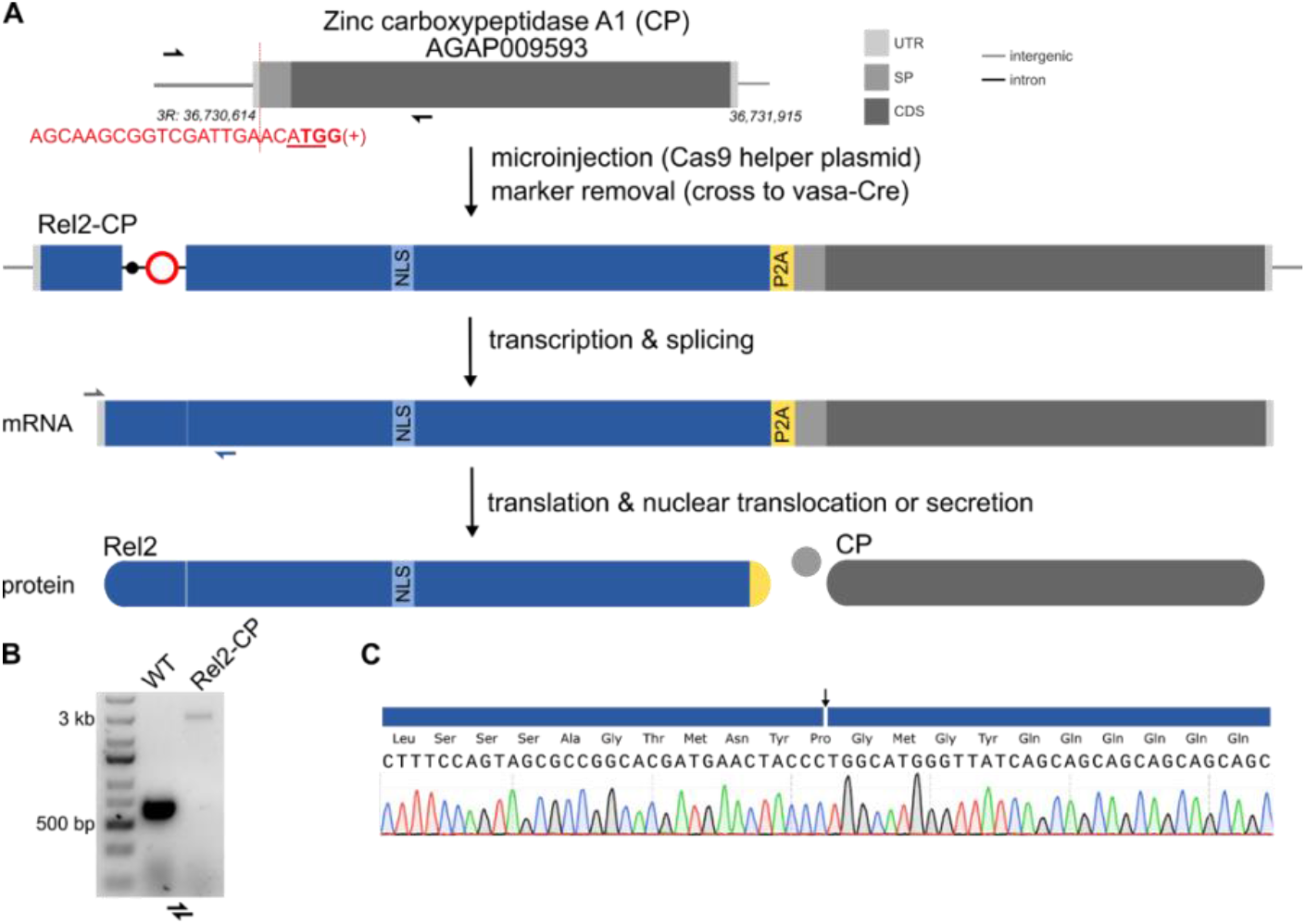
Transgenic line overexpressing REL2-S from within the CP locus. (**A**) Schematic illustrating the engineering of the CP host gene in REL2-CP mosquitoes, along with its transcription and translation processes. The gRNA target sequence (in red) encompasses the ATG start codon (underlined) that overlaps with the PAM motif (bold). Cre recombinase-mediated excision of the fluorescent marker module results in a markerless version of the transgene that contains only the gRNA module (red circle) and a loxP site (black dot) within the artificial intron. The NLS signal within REL2 is highlighted in light blue. Splicing of the artificial intron and autocleavage of the P2A peptide during translation lead to the production of two separate proteins: REL2-S, directed to the nucleus, and CP, secreted into the extracellular space. Half arrows denote the positions of PCR primers: black for genotyping in B and grey and blue for splicing confirmation in C. (**B**) PCR genotyping performed on genomic DNA extracted from a pool of 15 homozygous REL2-CP individuals and WT (KIL/G3) controls. (**C**) Sequencing of the amplicon obtained from RT-PCR on midgut cDNA demonstrates the precise splicing of the artificial intron.

Transgenesis of *An. gambiae* G3 was achieved through CRISPR/Cas9-mediated homology-directed repair, screening larvae for GFP expression in neuronal tissues. To establish a markerless version of the transgene, minimizing the presence of unnecessary exogenous sequences in this line, the GFP expression module within the artificial intron was flanked by loxP sites, enabling its Cre-mediated excision. Crossing the REL2^GFP^-CP line with a vasa-Cre line created in *An. gambiae* KIL strain background [12] led to the establishment of a homozygous markerless strain, REL2-CP, used in all subsequent experiments (**Figure 1B**). A KIL and G3 hybrid strain (KIL/G3) served as the WT control in these experiments. RNA extraction from midguts at 3h post-blood feeding (PBF), followed by cDNA splice site PCR and Sanger sequencing, confirmed correct splicing of the artificial intron (**Figure 1C**).

### REL2 localizes in the nuclei of posterior midgut epithelial cells

Since CP is expressed in the posterior midgut and upregulated after a bloodmeal [13], we anticipated a similar pattern of localization for the REL2-S effector. Despite the effector being fused to CP, which retains its secretion signal [14], the inclusion of a P2A cleavage peptide was expected to lead to a REL2-S protein that localizes in the nuclei due to its NLS. Spatiotemporal expression of REL2-S in the midgut tissue was visualized by indirect immunofluorescence (IFA), using an antibody targeting the residual aminoacids from the 2A peptide remaining at the C-terminus of REL2-S following ribosome skipping (**Figure 2**). Strong nuclear protein expression was detected in the posterior of blood-fed midguts at 6h and 20h PBF, with a slightly fainter signal at 25h PBF. No presence of REL2-S was observed in the cytoplasm, while some expression was detected in the anterior midgut of both blood-fed and non-blood-fed (NBF) midguts, largely limited to specific cell zones in the posterior part of the anterior midgut and the cardia that is also referred to as proventriculus.

**Figure 2.**
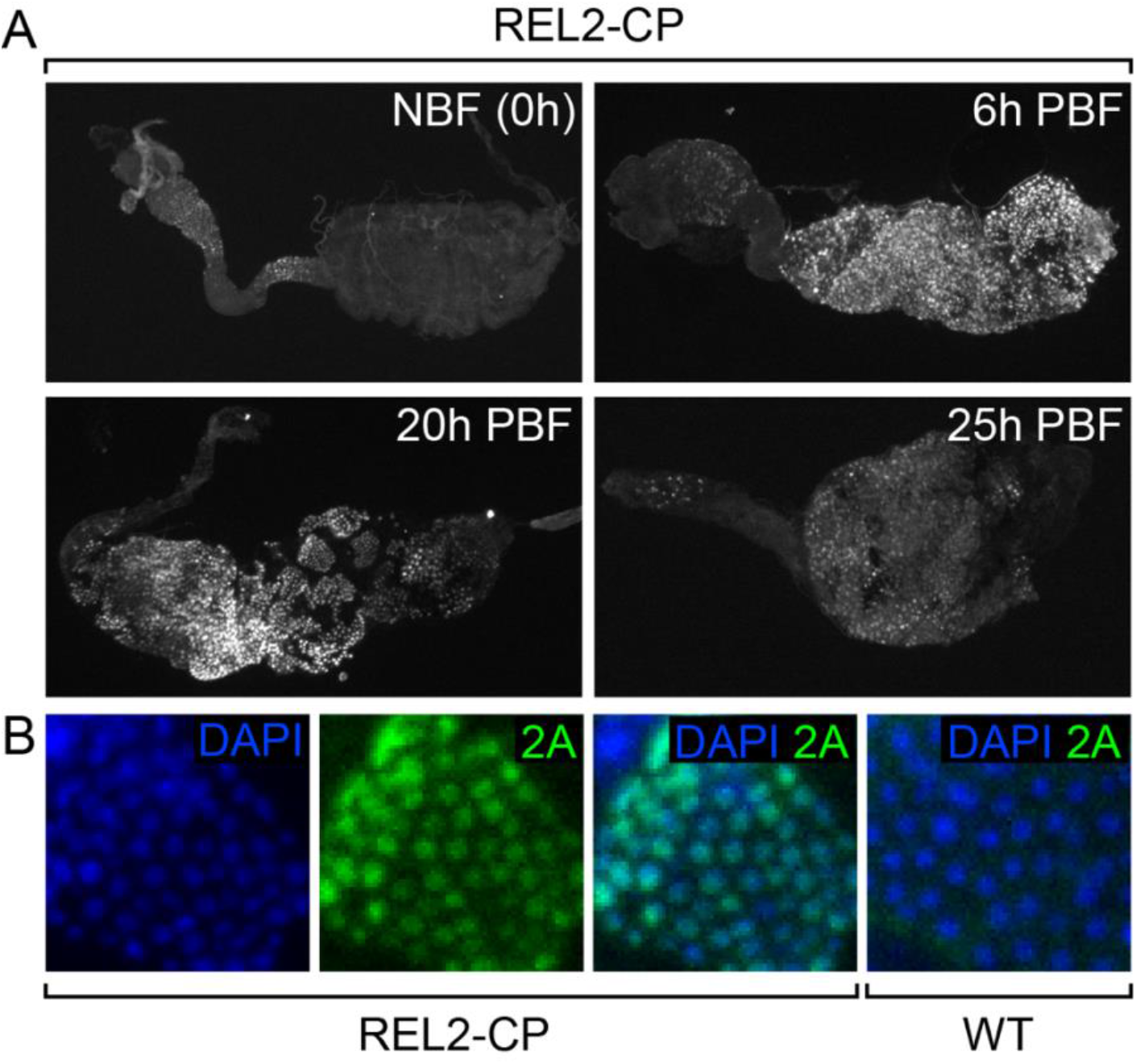
Spatiotemporal REL2-S protein expression. (**A**) Ectopic REL2-S protein expression is detected in the nuclei of posterior midgut cells via immunofluorescent staining with rabbit-anti-T2A at 6, 20 and 25 hours after bloodmeal (PBF). Expression in non-blood-fed (NBF) midguts sampled just prior to bloodfeeding (0h) was limited to the posterior part of the anterior midgut and the cardia. (**B**) Closeup view of REL2-S nuclear localization in the posterior midgut using the rabbit-anti-T2A antibody (2A). DNA is stained with DAPI.

### Phenotypic characterization of REL2-CP

We have previously described the generation and characterization of transgenic lines expressing the scorpine AMP from within the CP locus [11], and reported no statistically significant differences between those and the WT controls regarding fecundity or fertility, despite using the same female-biased and bloodmeal-inducible expression of the CP host gene. We assessed fecundity and fertility in females of the homozygous markerless REL2-CP line and compared them to the WT control. Egg output showed a significant reduction of approximately 50% (**Figure 3A**), but no statistically significant difference in larval hatching rate was noted (**Figure 3B**). The survival of males and blood-fed females was also reduced, with 49.1% of females deceased after 10 days, compared to 5.1% in the WT (**Figure 3C**).

**Figure 3.**
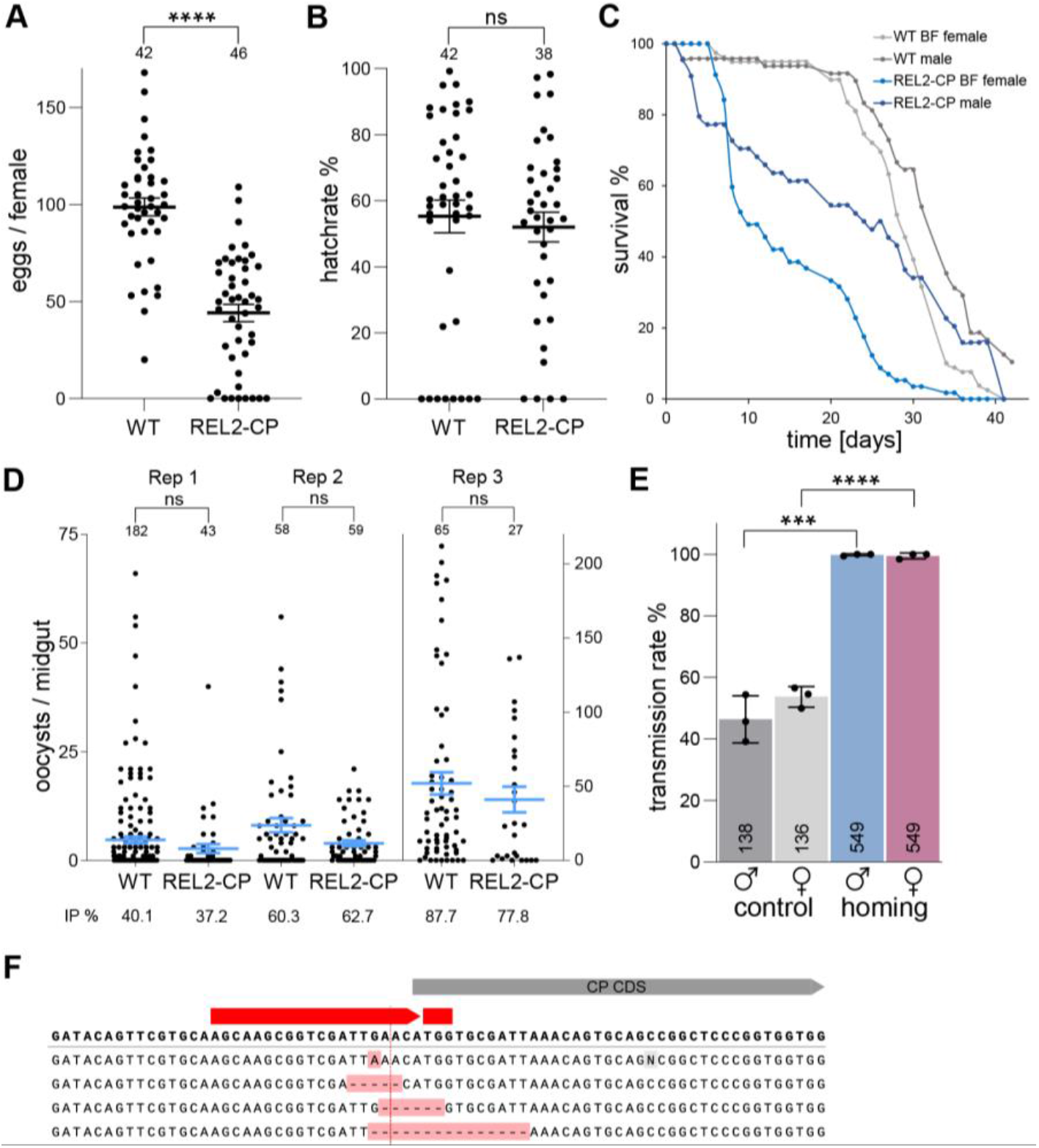
Phenotypic traits of the REL2-CP line. **(A)** Fecundity comparison between individual females of the homozygous REL2-CP and the WT strains. Statistical analyses here, as well as in (B) and (D, for infection intensity), were conducted using the Mann-Whitney test: ns, non-significant (i.e., p>0.05); ***, p<0.001. Mean and standard error of the mean (SEM) are presented, with the number of mosquitoes assayed noted above each dot plot. **(B)** Larval hatch rates derived from the eggs analyzed in (A). **(C)** Survival curves of males and females blood-fed on day 5 post-emergence. **(D)** *P. falciparum* oocyst intensity and prevalence in the midgut of REL2-CP compared to WT mosquitoes. Three biological replicates were performed, and mosquito numbers are indicated above the dot plots. For replicates 1 and 2, mosquitoes were starved for 48h PBF and dissected after 9 days, while mosquitoes in replicate 3 were starved for 72h PBF and dissected after 8 days. In the infection intensity graphs, the mean and SEM are shown. Infection prevalence is indicated below the infection intensity graph. **(E)** Transmission rates of the male and female REL2-CP transgene in control crosses to WT mosquitoes (grey bars) and in homing crosses to Cas9 expressing mosquitoes. Mean and SEM of 3 biological replicates is shown. Statistical significance was determined with unpaired t-test: ***, p<0.001; ****, p<0.0001. Total mosquito numbers are indicated within each column. **(F)** The 4 individuals that exclusively exhibited the WT band in the homing assay underwent PCR over the insertion site and the resulting sequences are shown. SNPs and indels are highlighted in red. The red vertical line indicates the cleavage site.

In infection experiments with *P. falciparum* NF54, both REL2-CP and WT were assessed for oocyst counts in the midgut. While the REL2-CP line exhibited a reduction in parasite infection intensity compared to the WT across three independent biological replicates, this difference did not reach statistical significance (**Figure 3D**).

We assessed the levels of germline gene drive in the markerless REL2-CP strain after crossing with a vasa-Cas9 line [10]. Trans-hemizygote offspring that derived from that cross were subsequently crossed with WT, and the transmission rate was assessed via multiplex PCR on larval progeny (**Figure 3E**). The results revealed high homing rates of 99.64% for males and 98.91% for females in trans-hemizygous REL2-CP (**Table S2**), comparable to those observed with Sco^GFP^-CP and Sco-CP [11]. Sanger sequencing of four individuals lacking the effector cassette showed that three exhibited deletions of variable length at the gRNA target site (**Figure 3F**).

### Knockout of the CP host gene reveals no significant fitness cost

To evaluate the potential contribution of the modified expression of the CP host gene to the observed phenotype, we generated a knockout strain (CP-KO) by inserting a cassette with a GFP expression marker 151bp downstream of the ATG of the CP gene (**Figure 4A**). This strain additionally contained a U6-gRNA module and was crossed to vasa-Cas9 to establish a homozygous colony, which exhibited no any obvious fitness issues during standard maintenance. CP knockout was confirmed via PCR on genomic DNA over the insertion site (**Figure 4B**), as well as RT-PCR on cDNA from 3h-blood-fed guts, which showed no expression of the CP gene (**Figure 4C**).

**Figure 4.**
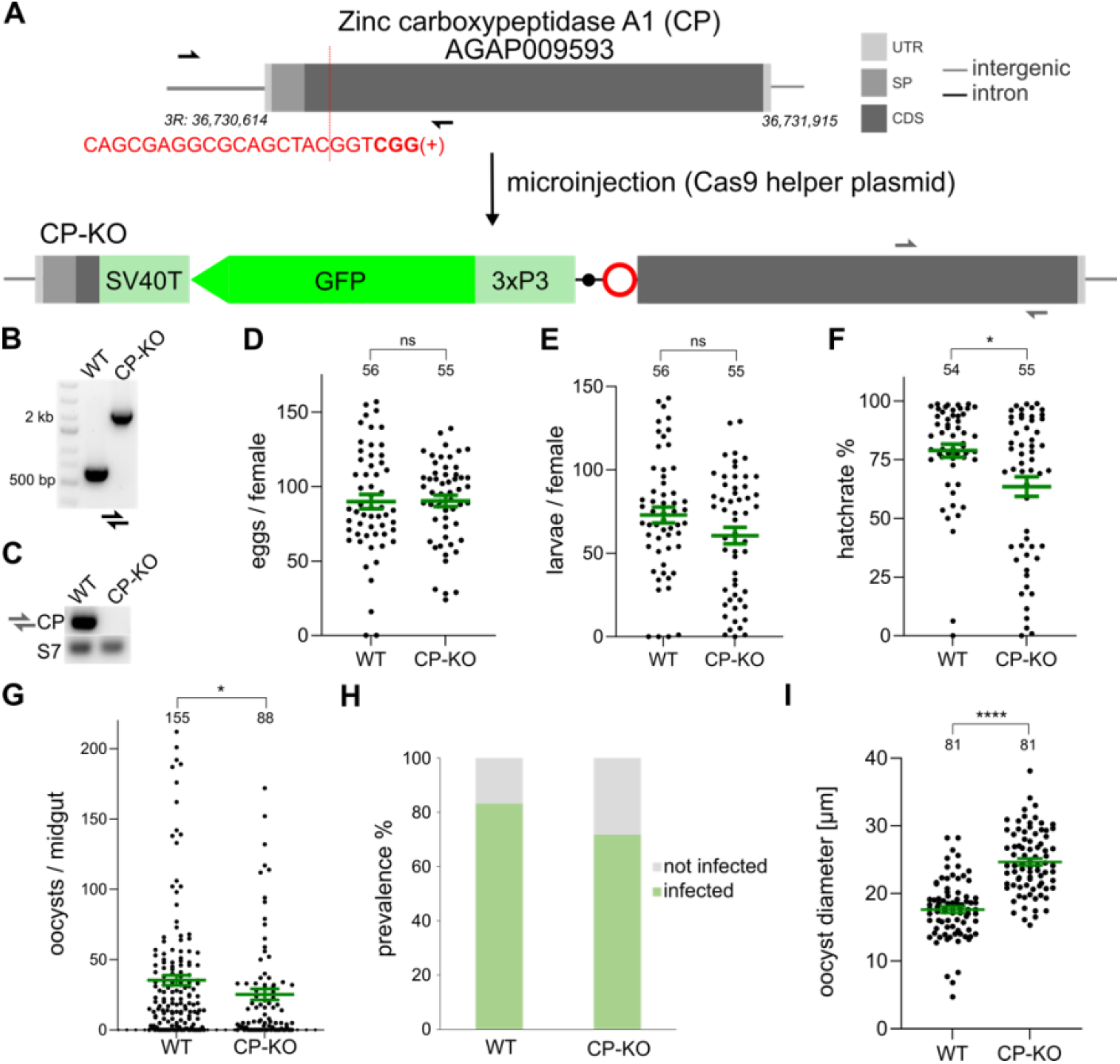
Knock-out of the CP host gene. **(A)** CRISPR/Cas9-mediated insertion of the knock-out construct into the endogenous locus of Carboxypeptidase (CP) with the gRNA target sequence (red) and the PAM motif (bold) indicated. The CP knock-out strain (CP-KO) contains a fluorescent eye marker module under the control of the synthetic 3xP3 promoter and the SV40 terminator, as well as an intronic guide RNA module (red circle) and one loxP site (black dot). Half arrows indicate primers for PCRs (black for genotyping in B, and grey for RT-PCR in C). **(B)** PCR genotyping on genomic DNA from a pool of 15 homozygous CP-KO individuals and WT control with primer-pairs spanning the entire locus (shown as black half-arrows in A). **(C)** RT-PCR on three-hour blood-fed midgut cDNA indicates knock-out of CP. The housekeeping gene S7 was run in parallel as reference. **(D)** Egg and **(E)** larvae of single females of the strain CP-KO compared to WT. **(F)** Larval hatch rates of the eggs and larvae shown in (D and E). p-values were calculated using the Mann-Whitney test. **(G)** *Plasmodium falciparum* oocyst intensity in the midgut of the transgenic strain CP-KO compared to WT. Females were left without sugar for 72 hours and dissected after 8 days. Data from three biological replicates were pooled and p-values were calculated via Mann-Whitney test. The number of midguts is indicated above, the mean and SEM are shown in green. **(H)** Infection prevalence (IP) of the three biological replicates in (G). **(I)** Oocyst diameter of the CP-KO compared to WT 8 days PBF analyzed using an unpaired t test, with the number of oocysts indicated above and the mean and SEM shown in green.

The number of eggs laid by single female CP-KO mosquitoes and the larval output was not significantly different compared to the KIL/G3 WT, but the larval hatching rate was found to be slightly reduced (**Figure 4D-F**). Infections with *P. falciparum* showed a small yet significant reduction in infection intensity for the CP-KO compared to the WT, but the reduction in infection prevalence was not significant (**Figure 4G-H**). We hypothesized that the knockout of the CP digestive enzyme might lead to a depletion of nutrients available for the parasite, which could cause a delay in growth and hence smaller oocyst size. However, the opposite was observed with oocyst diameter significantly bigger in the CP-KO than in the WT (**Figure 4I**). Overall, these findings suggest that CP is non-essential and possibly redundant, and that the fitness costs observed with the REL2-CP line are not due to the observed reduction in CP gene expression.

### Bloodmeal induced overexpression of REL2 causes broad transcriptional changes

The transcriptional activity of the CP locus is known to peak at about 3h PBF, and our observations had confirmed significant nuclear accumulation of recombinant REL2-S between 6h and 20h PBF. We chose these two timepoints to examine potential early and late changes in the midgut transcriptome of REL2-CP compared to WT female mosquitoes using RNA-seq (**Tables S3, S4, S5**). Midguts dissected just before bloodfeeding were used as a reference timepoint, designated as NBF. REL2-S overexpression in REL2-CP midguts was confirmed with log_2_ fold-changes of 3.3, 4.5, and 3.1 at NBF, 6h PBF and 20h PBF compared to WT controls, respectively. This suggests that there is transcriptional activity of the host CP locus prior to bloodfeeding and confirms maximum overexpression of REL2-S at 6 hours PBF (**Figure 5A**).

**Figure 5.**
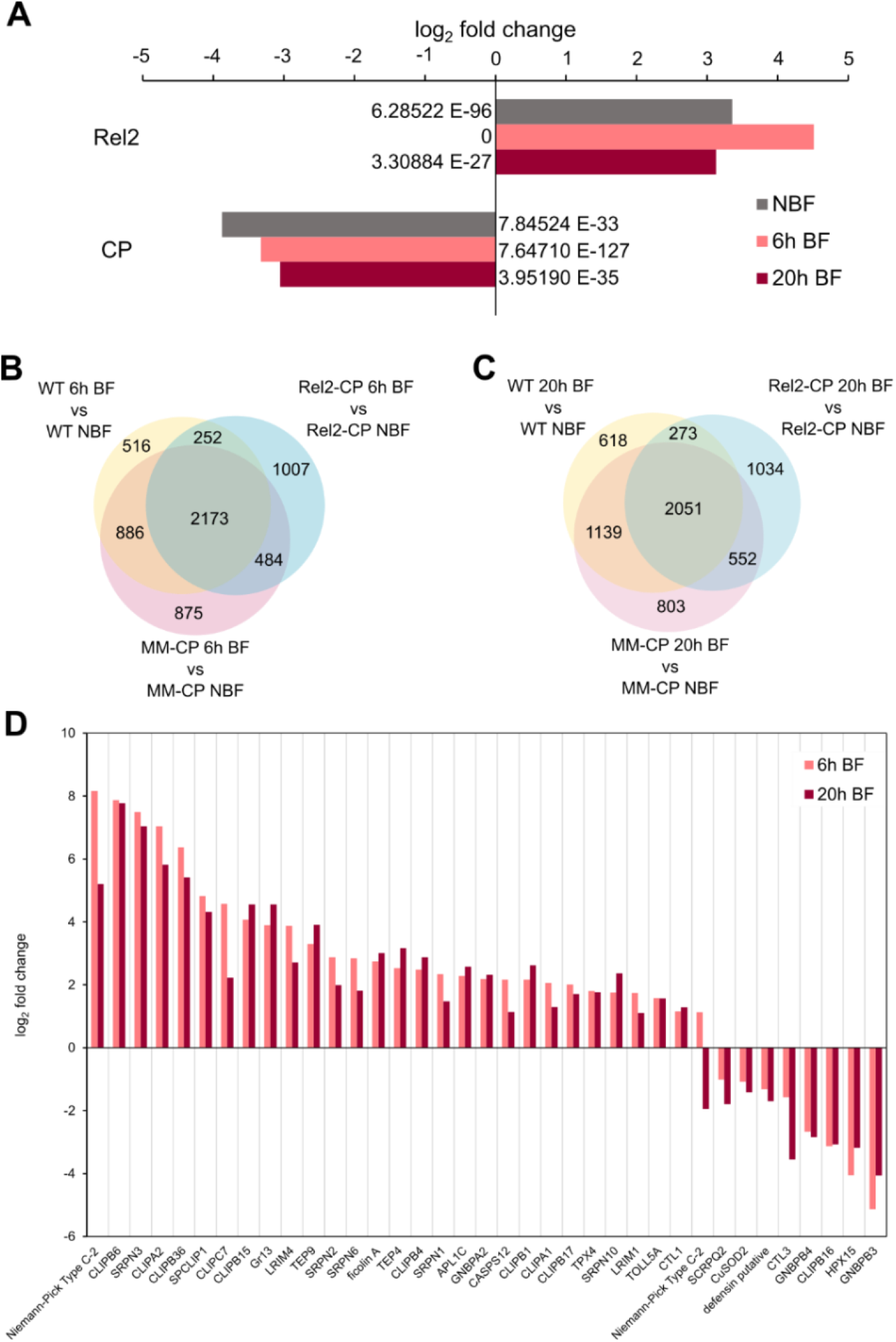
Transcriptome analysis of REL2-CP. **(A)** Log_2_ fold-change in mRNA expression of the effector REL2 and the host gene CP in the transgenic strain relative to WT. Adjusted p-values are indicated at the base of each column. **(B and C)** Venn diagrams of bloodmeal induced genes compared to WT and the previously published MM-CP strain [10] at 6 and 20 hours after feeding. **(D)** Immunity-related genes that are differentially expressed in REL2-CP at both blood-fed time points.

The REL2-S transgene, including the intron with the U6-gRNA module, spans 2.3 kb compared to only 860 bp in the MM-CP strain that has two short AMPs, Magainin 2 and Melittin, also integrated within the CP locus [10]. The RNA-seq data indicates that the CP locus is perturbed by this large integration, leading to significant reduction of CP host gene expression in the REL2-CP strain at all three timepoints examined (**Figure 5A**), whereas changes in CP levels were negligible in the MM-CP strain [10].

Expression analyses revealed the upregulation of a large number of genes upon feeding and significant midgut transcriptional changes between the REL2-CP and WT mosquitoes (**Figure S1**). Next, we co-analyzed the RNA-seq data with those published previously for the MM-CP line [10]. Across all three timepoints, over 90% of genes were co-expressed in WT, REL2-CP, and MM-CP (**Figure S2**). However, 1.97 and 6.75 times more genes were found to be uniquely expressed in the REL2-CP compared to the other two lines (**Figure S2**). In comparison to WT, 1.95 and 1.67 more genes were induced by a bloodmeal in the REL2-CP at 6h and 20h PBF, respectively (**Figure 5B-C**).

Comparing the data from 6h PBF midguts to a previously published microarray study of the *An. stephensi* CpREL2_15_ transgenic line [7], we found that out of the 191 genes upregulated in CpREL2_15_ (p-value < 0.05, log_2_ fold-change > 0.75) only 14 were among the 714 upregulated in our RNA-seq data (adjusted p-value < 0.05, log2 fold-change > 1). When we applied a more stringent threshold for log_2_ fold-change in our data, due to generally higher values in the RNA-seq compared to the microarray study, only 3 genes overlapped between the 115 and 743 genes downregulated in the *An. stephensi* CpREL2_15_ and REL2-CP lines, respectively.

### Transcriptional changes in immunity, metabolism and oogenesis related genes

GO enrichment analysis of REL2-CP compared to WT DEGs at 6h PBF revealed 14 enriched GO-terms exclusively in upregulated genes (**Table S6**). These terms included GTP, guanyl, and purine nucleoside/nucleotide binding, as well as catalytic activities acting on RNA and tRNAs, transferase, and ligase activities, indicating heightened transcriptional processing in the REL2-CP midguts. Conversely, downregulated genes exhibited 49 GO-terms, mainly related to electron transfer, energy, and respiration. At the 20h PBF timepoint, upregulated genes were significantly enriched in GO-terms related to translation initiation, amino acid activation, protein folding, protein glycosylation, vesicle-mediated transport, protein localization, unfolded protein response and proteolysis, indicating increased translation and subsequent processes aligned with increased protein synthesis and processing.

No significant GO-term enrichment was observed for immunity-related genes at either timepoint PBF. However, when comparing the DEGs to a curated list of 288 genes involved in immunity, 42 and 45 genes were upregulated at 6h and 20h PBF, respectively, with 30 of them overlapping between both timepoints (**Table S7**). Among them were 14 CLIP-domain serine proteases (CLIPs), 6 serpins (SRPNs), 3 Niemann-Pick type proteins, 4 C-type lectins (CTLs), and 3 thioester-containing proteins (TEPs). Additionally, 16 and 25 of the DEGs downregulated at 6h and 20h PBF, respectively, were related to immunity, including 3 Class B Scavenger Receptors (SCRBs), 3 heme peroxidases (HPXs), and 2 prophenoloxidases (PPOs). Several members of the leucine-rich repeat immune protein (LRIM) family, including the complement-like pathway factors LRIM1 and APL1C, were also upregulated. Surprising, none of the AMPs were transcriptionally affected.

Next, we investigated which genes might be responsible for the observed reduction in egg output in REL2-CP. For this, we considered genes with GO terms related to oogenesis and identified 11 DEGs, primarily heme peroxidases and cytochrome P450s (**Table S8**). Notably, Nanos, maternally deposited into the egg [15], was 3.6-fold downregulated at 6h PBF. Additionally, we observed a 3.2-fold downregulation at the same timepoint of the ovo orthologue (AGAP000114), a Zinc finger transcription factor in *Drosophila* with a dual role as a repressor or activator, resulting in female sterility when mutated.

We also examined genes associated with juvenile hormone (JH), which is crucial for egg production in insects [16]. Among the 12 DEGs, 6 were upregulated at 6h PBF (**Table S9**), including JH esterase and epoxide hydrolases. JH esterase was also reported to be induced in transgenic *Aedes* overexpressing REL2 under the Vg promoter [5]. Since only three *Anopheles* genes are associated with GO-terms related to lifespan and aging, establishing a direct linkage between DEGs and the observed decrease in adult survival was not possible.

Finally, comparing the DEGs to a list of 75 genes involved in metabolism upon bloodfeeding [17], we found a total of 26 affected genes, with 14 being upregulated and 12 being downregulated (**Table S10**), indicating some influence of REL2-S overexpression on bloodmeal digestion.

### Molecular signatures in the upstream regions of target genes

To gain insights into the regulatory mechanisms underlying the differential gene expression observed upon bloodmeal-induced overexpression of REL2-S, we conducted an analysis of the upstream regions of DEGs. Initially, we examined transcription factors within *Drosophila* motif databases with binding preferences similar to REL2, which could potentially act as co-activators or co-repressors. We identified *Drosophila* Dorsal, but its *An. gambiae* orthologue REL1 showed no differential expression. Schnurri (shn), known to function as both a co-activator and co-repressor involved in midgut development, and co-activator Akirin [1] were also identified. However, their orthologues were also not among the DEGs.

Previous *de novo* motif discovery using open chromatin profiling by FAIRE-seq on *An. gambiae*-derived immune-responsive cells had revealed sites for STAT, lola, and Deaf1-type transcription factors [18]. Nonetheless, the orthologues of these genes were either downregulated or not differentially expressed.

The transcription factors Caudal (Cad), Nubbin (Pdm1), zinc finger homeobox protein (Zfh1/2), and Polybromo (Bap180) are recognized for their role in modulating Imd pathway activity, leading to the repression of AMP expression [19]. Silencing Cad through RNAi was shown to increase CEC1 expression in *An. gambiae* midguts [20]. However, the *An. gambiae* orthologues of these genes were either downregulated or not differentially expressed.

To distinguish between direct and indirect target genes, despite the absence of a positional weight matrix (PWM) for *An. gambiae* REL2, we leveraged the highly conserved residues binding to DNA, allowing us to use a Drosophila Y2H matrix [21] to screen the 10kb upstream regions of DEGs. Of the bloodmeal-induced DEGs, 29.97% of upregulated (232/774) and 23.41% of downregulated (291/1243) genes had no assigned REL2 binding site, suggesting they might be indirect targets or affected by compensatory mechanisms. Conversely, genes with 3 or more non-overlapping REL2 sites within 2.5 kb upstream from the ATG (excluding exons from neighboring genes) were considered likely direct targets. Ten genes were identified within the upregulated set, including neuropeptide pyrokinin capa-like (CAPA), odorant-binding protein 10 (OBP10), and JAK/STAT pathway cytokine unpaired 3 variant A (UPD3A) (**Table S11**). Three genes were identified within the downregulated set, namely CEC1, a cystinosin, and a leucine-rich repeats and immunoglobulin-like domains protein 3 (LRIG3). All those upstream regions contained 3 to 4 REL2 transcription factor binding sites (TFBS).

We then focused on AMPs that are thought to be direct targets of immune signaling pathways, including cecropins (CEC1-3), defensins (DEF1-5) and gambicin (GAM1). While CEC2 was not among DEGs and exhibits no recognizable REL2 binding sites, CEC1 and CEC3 share 5 REL2 binding sites in their intergenic region and were both downregulated upon blood feeding. Similarly, DEF2-5 that showed no recognizable REL2 binding sites were either downregulated or not among DEGs, while DEF1 with 4 potential REL2 binding sites was also downregulated PBF. GAM1, which contains one REL2 binding site in its upstream region, was also downregulated 20h PBF. These data together implied that overexpression of REL2 may have the opposite effect on its targets by directly acting as a repressor or inducing the expression of other transcriptional repressors.

To investigate the latter, we initially considered genes associated with GO-terms related to repression (GO:0001217 and GO:0070491). Among the 17 genes with these GO-terms, seven were differentially expressed. However, only AGAP012346, encoding an unspecified product, was upregulated after a bloodmeal and lacked an annotated orthologue in *Drosophila*. Next, we considered GO-terms associated with transcription (GO:0006355, GO:0003700, GO:0006351, GO:0008134) and identified 49 upregulated and 57 downregulated genes among the DEGs (**Table S12**). Notably, several putative repressors were upregulated, including the orthologues of single-minded (AGAP000773), a master regulator of nervous system development [22], snail (AGAP008274), involved in mesoderm development, SOX1/3/14/21 (SOX group B, AGAP010919), with dual roles as activators or repressors in vertebrates [23], hairy and enhancer of split (AGAP012342), participating in the control of cell fate choice, and POU domain transcription factor (class 6, AGAP012345), serving as both an activator and repressor [24].

We expanded our screen to all potential transcription factors. Out of 66 upregulated putative transcription factors with *Drosophila* orthologues, 18 had available PWMs. We scrutinized the 3.5 kb regions upstream of the ATG of *An. gambiae* CEC1, DEF1, and GAM1 for potential binding sites of these transcription factors using Drosophila PWMs. In all three genes, we identified a TALE homeobox transcription factor (CG11617) binding site. Additionally, binding sites for Deadpan (dpn), Clock-Cycle (clk-cyc) heterodimer, and E(spl)mβ-HLH (Enhancer of split mβ, helix-loop-helix) were found upstream of DEF1 and GAM1. Dpn and E(spl)mβ-HLH act as transcriptional repressors for genes requiring a basic helix-loop-helix (bHLH) protein for transcription, while the Clk-cyc activator complex regulates circadian rhythms. In the regulatory region of GAM1, hits for Snail (sna), Sugarbabe (sug), Suppressor of Hairless (Su(H)), Single-minded (sim), and Hormone receptor 3 (Hr3) were also observed. Sug represses a set of lipase genes involved in fat catabolism, Su(H) activates transcription of Notch target genes when associated with Notch protein but represses when not, and Hr3 is induced directly by 20-hydroxyecdysone. Additionally, binding sites for E(spl)m3-HLH and E(spl)m7-HLH were detected, often overlapping with Dpn.

Finally, putative C/EBP (CCAAT/enhancer binding protein) binding sites have been previously found to be closely associated with REL2 binding sites in DEF1 and the *Ae. aegypti* DefA1, and are believed to contribute to target gene upregulation when overlapping with and downregulation when found near the REL2 binding sites [25, 26]. Since C/EBP (AGAP011096) was downregulated in our RNA-seq data, we hypothesize that an imbalance in the REL2 to C/EBP ratio may also contribute to the lack of AMP activation.

## Discussion

Overexpressing the NF-κB-like transcription factor REL2 in *Anopheles* mosquitoes has long been seen as a promising strategy for genetically controlling malaria transmission. Drawing on the well-established Imd pathway, which regulates the expression of AMPs and other immune effectors in the *An. gambiae* midgut, we adopted a previously successful approach of expressing REL2-S under the control of the CP promoter [6-8] within our IGD framework by integrating REL2-S within the CP locus [9, 11]. In this configuration, the trangenic REL2-CP locus contains all the necessary elements to drive population-wide modification in the presence of a Cas9 source. Our study confirmed robust upregulation of REL2 in the midgut of transgenic mosquitoes, with the protein localizing in the nuclei of epithelial cells in the posterior midgut after a bloodmeal. This induction causes significant transcriptional changes and results in a marked decrease in both oviposition and lifespan. However, the expected reduction in *P. falciparum* infection was minimal, likely due to the failure of this approach to induce the expression of AMPs and other immune effectors.

CP is considered important for blood digestion. We previously demonstrated that CP can be a suitable host gene for antiparasitic effectors, and its expression was only minimally affected when short AMPs were inserted within this gene [10, 11]. However, when the relatively long REL2-S was inserted at the start of the CP CDS, the expression of CP was significantly downregulated. To assess the impact of this reduction on the REL2-CP phenotype, we generated and characterized a transgenic CP knockout strain. In contrast to our expectations, the results revealed that CP is non-essential and possibly redundant, with no observable effects on mosquito fecundity or fertility. Consequently, the considerable fitness cost observed in the REL2-CP strain is attributed to the expression of REL2 and not the decreased CP protein levels.

The principal concept of IGD revolves around the expression of the effector gene from within host genes, capitalizing on the natural expression mechanisms of those genes. Typically, a host gene with robust tissue-specific and spatial expression is chosen. However, the selection of host genes exhibiting strong expression might not always be the optimal choice, especially when dealing with effector transcription factors, like REL2. Our RNA-seq data uncovered numerous responses to the effector component, likely stemming from indirect effects, compensatory mechanisms, microbiota interactions, host locus associations, or secondary alterations of gene expression. This caused changes in genes linked to blood digestion, juvenile hormone metabolism and the gonotrophic cycle, leading to defects in oogenesis and providing a plausible explanation for the observed decrease in female fecundity. Such substantial transcriptional shifts can have serious implications for the overall mosquito fitness leading to the observed increase in mortality rates.

Indeed, a quarter of the bloodmeal induced DEGs in REL2-CP mosquitoes are unlikely to be directly regulated by REL2, as no putative REL2 binding sites were detected in their upstream regions. Only 10 of the upregulated DEGs exhibited 3-4 REL2 binding sites, indicating them as potential primary targets of the REL2 transcription factor. Intriguingly, among downregulated DEGs, we identified 3 genes with a notable clustering of REL2 binding sites in their promoter region: CEC1, AGAP004115, a homolog of the lysosome cystine transporter cystinosin, and AGAP009688, a homolog of LRIG3.

In our previous study, we demonstrated that overexpressing exogenous AMPs in the posterior mosquito midgut post-bloodmeal could impact *P. falciparum* infection, holding promise for potential malaria transmission blockade [10]. Consequently, the intended outcome of REL2 overexpression was the transcriptional activation of Imd pathway-controlled AMPs in the posterior midgut. Although the minimal effect on infection and the lack of AMP upregulation in the REL2-CP mosquitoes was initially surprising, a detailed analysis of the upstream regions of target AMPs provided some clarity.

CEC2 and DEF2-5, all lacking REL2 binding sites in their regulatory regions, as well as GAM1, with only a single REL2 binding site, did not show any response. However, DEF1, CEC1, and CEC3, each housing 4-5 REL2 binding sites in their regulatory regions, were downregulated post-bloodfeeding.

These findings suggest that AMPs controlled by the Imd pathway undergo transcriptional repression in the REL2 overexpression background, and that REL2 expression and nuclear translocation alone are insufficient to activate their expression. Instead, REL2 overexpression appears to induce the upregulation of many transcriptional repressors, some of which may act as a feedback mechanism to control or dampen the Imd pathway activation. Given that the microbiota harbored in the posterior midgut are critical for the mosquito fitness, this may be a tolerance mechanism that ensures microbiota persistence by suppressing the overactivation of immune responses.

The *An. gambiae* orthologues of established nuclear repressors in the Imd pathway, such as Caudal, Nubbin, Zfh1, and Bap180, were found to be either downregulated or not differentially expressed in our study. However, our investigation revealed several known repressors not previously associated with immune repression, which were upregulated in REL2-CP mosquitoes post-bloodmeal. These included Single-minded, Snail, SOX1/3/14/21, a POU domain transcription factor, Hairy, and enhancer of split. Importantly, the regulatory regions of DEF1 and GAM1 do harbor binding sites for some of these factors suggesting a plausible link between the lack of activation and, indeed, the suppression of AMPs and the simultaneous upregulation of these potential repressors in REL2-CP mosquitoes post-bloodmeal.

## Material & Methods

### Plasmid construction

The short form of *An. gambiae* REL2 (AGAP006747) was amplified from the pBac-EGFP[AgCp-REL2-TryT] plasmid [6], and the artificial intron was inserted into the REL2-S CDS at position 227 between the CCC and TG sequences. The intron including the gRNA and the backbone including both CP homology arms were amplified from pD-Mag-Mel-CP [10]. Those 4 fragments were fused via Gibson assembly to yield the donor plasmid pD-REL2-CP. The *Drosophila* splice predictor [27] was used to predict the splicing probability of the effector cassette.

The gRNA for the CP-KO was introduced into the intermediate plasmid pI-Scorpine [11] via Golden Gate cloning with BbsI, and the marker-gRNA-module was PCR-amplified with primers 599-SV40-KO-F and 600-scaff-KO-R. The 5’ and 3’ homology arms were amplified from G3 gDNA and fused with the backbone and the marker-gRNA-fragment to yield the donor plasmid pD-CP-KO. For primers see **Supplementary Table S1** and for the donor plasmids see **Supplementary Files 1 and 2**.

### Microinjection and establishment of markerless versions

*An. gambiae* G3 eggs were injected with the donor plasmid pD-REL2-CP and the Cas9 helper plasmid p155 [28]. Eighteen transient survivors and 23 F1 transgenics were obtained, and the REL2^GFP^-CP line was established from a founder cage with 2 males. Correct insertion was confirmed by Sanger-sequencing on 9 F2 individuals. Transgenic mosquitoes were outcrossed to G3 WT over two generations before crossing to the vasa-Cre strain [12]. Larval offspring were screened for GFP and DsRed and siblings were mated. The progeny was screened against GFP and DsRed, and genotyped on single pupae exuviate with primers 99-CP-locus-F and 100-CP-locus-R to identify homozygotes, and a cage was set up with 1 female and 7 males. To boost the colony, the offspring were crossed to the vasa-Cas9 line (E. Marois, unpublished), progeny was sib-mated and maintained over Cas9 without screening for 6 generations. Subsequently, the larvae were screened against Cas9, control-genotyping was performed on a subset, and no WT was detected.

*An. gambiae* KIL/G3 eggs were injected with the donor plasmid pD-CP-KO and the Cas9 helper plasmid p155 [28]. 18 transient survivors and 10 F1 transgenics were obtained and pD-CP-KO was established from a founder cage with 3 males, which were confirmed by Sanger-sequencing. Transgenics were outcrossed to KIL/G3 WT over two generations before crossing to vasa-Cas9 (E. Marois, unpublished). Larval offspring were screened for GFP in the CFP channel and for YFP in the mKO/mOrange channel and siblings mated. The progeny was screened for GFP and against YFP.

Crosses of 2 females with 2 males were set-up in cups, blood-fed, allowed to lay eggs, and the parents were genotyped with primers 99-CP-locus-F and 100-CP-locus-R to confirm homozygosity.

### RNA extraction, RT-PCR and RNA-seq

REL2-CP and the WT control were fed with human blood and 15 midguts were dissected. After lysis in Trizol and homogenization with 2.8mm ceramic beads (CK28R, Precellys) for 30s at 6,800rpm in a Precellys 24 homogenizer (Bertin), RNA was extracted with the Direct-zol RNA Mini-prep kit (Zymo Research) including on-column DNase treatment. For splicing confirmation, RNA from 14 3h PBF guts was transcribed into cDNA with the iScript gDNA Clear cDNA Synthesis Kit (Bio-Rad) and RT-PCR was performed with RedTaq Polymerase (VWR) using primers 340-CP-Taq-F and 264-qRel-R2. Four biological RNA replicates before and at 6 hours and 20 hours after bloodfeed were subjected to RNA-seq via Illumina platform (Novogene) in parallel with the published MM-CP, as described previously [10]. Carcasses were collected and genotyped with primers 531-CP-multi-R and 532-CP-multi-F to confirm integrity of the mosquitoes used for the experiments.

CP-KO RNA was extracted from 15 3h PBF guts with the Direct-zol RNA Mini-prep kit (Zymo Research) including on-column DNase treatment and transcribed into cDNA with the qScript cDNA Synthesis Kit (Quantabio). RT-PCR was performed with RedTaq Polymerase (VWR) using primers 154-CP-probe-F & 155-CP-probe-R, as well as 447-S7-F & 448-S7-R as housekeeping reference gene.

### RNA-seq data analysis

Enrichment analysis was performed with ClusterProfiler v3.8.1 [29] (padj < 0.05) and subsequently redundant GO terms were removed via Revigo with default settings [30]. Only Genes with an adjusted p-value < 0.05 and log_2_ fold-change > 1 were considered for further analysis. For the comparison to the published microarray of *A*.*s*. CpREL2_15_, data were taken from the supplementary table S3 [7], and for the cases of several contigs matching to the same AGAP number, the more extreme value was taken. For the analysis of DEG responsible for reduced egg laying, the terms egg, oogenesis, oocyte, ovarian and nurse cells were used to mine Amigo2 [31] for GO-terms, which were then used as input for g:Profiler [32] to extract *Anopheles gambiae* genes. For the comparison to the published list of genes involved in metabolism after bloodmeal [17], EST-IDs from the supplementary table S1 were converted to AGAP numbers. 7 EST-IDs had no AGAP number associated with it, 3 EST-IDs had two AGAP numbers associated with it, and 4 duplicates were removed, leaving 75 genes out of the initial 84 EST-IDs.

### Immunofluorescence assays (IFA)

Midguts were dissected, opened to remove the blood bolus, fixed in 4% paraformaldehyde and blocked in 1% (w/v) BSA, 0.2% (v/v) Triton-X. Tissues were incubated with polyclonal rabbit-anti-T2A (1:500, crb2005269, Discovery Antibodies) overnight and subsequently incubated with goat-anti-rabbit-IgG-Alexa Fluor 488 (1:1000, A11034, Invitrogen), stained with DAPI, mounted in ProLong Gold antifade reagent (Invitrogen) and imaged on EVOS FL (Invitrogen) with a 4x objective.

### Phenotypic assays

For the fecundity and fertility assay, 15 females were fed with cow-blood, transferred to cups singly and data from three biological replicates were pooled. Spermatheca were dissected from females that failed to give eggs and/or larvae in order to exclude them from the analysis if no sperm was detected. Pooled data did not show a Gaussian distribution according to Shapiro-Wilk and the strains were compared via Mann-Whitney test. For the survival assay, 50 male or female pupae, respectively, were allowed to hatch in a cage. Females were blood-fed 5 days after emergence and unfed individuals were removed.

Infections of mosquitoes with mature *Plasmodium falciparum* NF54 gametocyte cultures (2-6% gametocytaemia) were performed as described previously using the streamlined Standard Membrane Feeding Assay [33]. After infectious bloodmeal, females were starved without sugar for 48h (REL2-CP replicate 1 and 2) and 72h (REL2-CP replicate 3 and all replicates of CP-KO). Midgut dissections were performed on day 8 (REL2-CP replicate 3 and all replicates of CP-KO) and day 9 (REL2-CP replicate 1 and 2) after infection.

### Homing analysis

At least 56 homozygous REL2-CP individuals were crossed to the vasa-Cas9 strain carrying the 3xP3-YFP marker and the progeny were screened for the presence of yellow fluorescence (Table S2). In parallel, control-crosses to WT instead of Cas9 were prepared. Transhemizygotes were then sexed and crossed to WT, and gDNA was isolated from the offspring L2-L3 larvae according to the storage & dilution protocol of the Phire Tissue Direct PCR Kit (Thermo Scientific). Multiplex PCR was performed with primers 530-P2A-multi-F, 531-CP-multi-R and 532-CP-multi-F, yielding a 477bp band if the construct is present and a WT band of 670bp as control. Two 96-well-plates per parent (paternal or maternal transhemizygotes) and replicate were analysed and four negative controls were included on each plate, three with the dilution buffer and one with water. From the control-crosses, 46 offspring per parent were analysed for each replicate. The homing rate was calculated as (n*0.5 – E_neg_) / (n*0.5) *100, assuming a Mendelian distribution of 50% as baseline, with E_neg_ being the individuals negative for the effector and n being the number of larvae analysed excluding the PCRs that did not work. The four individuals that did not produce the band for the construct were subjected to PCR with primers 531-CP-multi-R and 532-CP-multi-F and the product was sequenced.

Deconvolution of the Sanger-chromatograms was done with CRISP-ID [34].

### Transcription factor binding site analysis

FIMO was used to scan sets of regulatory regions for individual matches to the motifs of interest [35]. TOMTOM was used to compare the REL2 motif against other known motifs within the combined Drosophila databases [36].

## Acknowledgments

We are grateful to Yuemei Dong and George Dimopoulos for the pBac-EGFP[AgCp-REL2-TryT] plasmid and Eric Marois for the vasa-Cre and vasa-Cas9 lines. The work was funded by the Bill and Melinda Gates Foundation grant OPP1158151 to N.W. and G.K.C.

## Author contributions

AH: Conceptualization, Formal analysis, Investigation, Investigation, Supervision, Vizualization, Writing – original draft; PC: Investigation, Methodology; GDC: Investigation, Methodology; MGI: Methodology; TH: Investigation; JAC: Data curation, Resources; GSS: Investigation; HN: Investigation; NW: Conceptualization, Funding acquisition, Project administration, Supervision, Writing – review & editing; GKC Conceptualization, Funding acquisition, Project administration, Supervision, Vizualization, Writing – review & editing.

## Competing interests

The authors declare that no competing interests exist.

## Supplementary figures

**Figure S1. RNA-seq heatmap after clustering of replicates**.

REL2-CP (Rel) and wild-type (WT) at non-blood-fed (N), 6 hours (6) and 20 hours (20) after bloodmeal from four biological replicates (1-4).

**Figure S2. Co-expressed genes**.

Comparison with the previously published MM-CP strain.

**Figure S3. Alignment of the *Drosophila* Relish and *An. gambiae* REL2 sequences**.

Amino acids contacting the DNA are highlighted in green or orange depending on whether they coincide in *An. gambiae* and *Drosophila* Rel, or not, respectively.

## Supplementary tables

Table S1. Primers used in this study.

Table S2. Transmission rates and homing rates.

Table S3. RNA-seq data NBF.

Table S4. RNA-seq data 6h BF.

Table S5. RNA-seq data 20h BF.

Table S6. Enriched GO-terms.

Table S7. DEG related to immunity.

Table S8. DEG related to oogenesis.

Table S9. DEG related to juvenile hormone.

Table S10. DEG related to metabolism.

Table S11. DEG related to transcription.

Table S12 DEG with at least 3 REL2 TFBS in their regulatory region.

## Supplementary files

Supplementary File 1. GenBank-DNA-files of the donor-plasmid pD-REL2-CP Supplementary File 2. GenBank-DNA-files of the donor-plasmid pD-CP-KO.

## Notes

### Competing Interest Statement

The authors have declared no competing interest.

